# Quadratic Mutual Information estimation of mouse dLGN receptive fields reveals asymmetry between ON and OFF visual pathways

**DOI:** 10.1101/2020.12.03.409342

**Authors:** Zhiguang Mu, Konstantin Nikolic, Simon R. Schultz

## Abstract

The longstanding theory of “parallel processing” predicts that, except for a sign reversal, ON and OFF cells are driven by a similar pre-synaptic circuit and have similar visual field coverage, direction/orientation selectivity, visual acuity and other functional properties. However, recent experimental data challenges this view. Here we present an information theory based receptive field (RF) estimation method - quadratic mutual information (QMI) - applied to multi-electrode array electrophysiological recordings from the mouse dorsal lateral geniculate nucleus (dLGN). This estimation method provides more accurate RF estimates than the commonly used Spike-Triggered Average (STA) method, particularly in the presence of spatially correlated inputs. This improved efficiency allowed a larger number of RFs (285 vs 189 cells) to be extracted from a previously published dataset. Fitting a spatial-temporal Difference-of-Gaussians (ST-DoG) model to the RFs revealed that while the structural RF properties of ON and OFF cells are largely symmetric, there were some asymmetries apparent in the functional properties of ON and OFF visual processing streams - with OFF cells preferring higher spatial and temporal frequencies on average, and showing a greater degree of orientation selectivity.

## I. INTRODUCTION

Consider two image patches of identical size but opposite contrast. A light patch on a dark background is perceived differently to a dark patch on a light background: Helmholtz termed this illusion the “irradiation illusion” [1]. It arises partly because of the scattering effects of light, however Galilei suggested that this may also indicate a difference in spatial resolution [2]. Schiller and colleagues [3] found that 2-amino-4-phosphonobutyrate (APB) abolishes ON response in retinal ganglion cells (RGCs), the LGN and visual cortex, but has no effect on the center surround antagonism of OFF cells or the orientation and direction selectivity in the cortex. These and related findings suggest that the ON and OFF pathways remain largely separate though the LGN and cortex [4], opening up the potential for the separate pathways to be differently adapted to specific information processing demands.

Asymmetry between ON and OFF blue sensitivity-cones of retina was observed as early as in the 70s by de Monasterio [5]. Other asymmetric properties such as the size of the receptive field of ON and OFF visual pathway have been observed in drosophila [6], rodents [7], tree shrews [8], leporids [9], cats [10] and primates [11], either in retina or visual cortex. The fact that this feature commonly existed among a diverse variety of species illustrates it may carry great importance in the functional role as a result of early development in the evolution [12]. In addition, the asymmetry in ON and OFF visual pathway has been linked with their physical properties, functions and information processing capabilities, this includes quantity [13], morphology & receptive field size [14], contrast sensitivity [15], direction tuning [9], motion detection [6], depth perception [4] and velocity estimation [16].

Teasing apart potential information processing asymmetries between ON and OFF receptive fields requires, as a first step, accurate estimation of RF structure. In many previous studies, spike-triggered average (STA) estimates have typically been used. In many cases, the statistical inefficiency of this approach has required averaging across time to produce a purely spatial RF; as well as resulting in a decreased number of apparently responsive units (in cases where there are temporally separated ON and OFF components that nearly average out over time), it can disguise ON-OFF asymmetries. There is thus the need for improved RF estimators. For a linear RF model with Gaussian noise, minimising the mean squared error corresponds to a maximum likelihood estimation (MLE) or a maximum a posterior estimation (MAP) with uniform prior. If in addition, the stimulus is spatial-temporally decorrelated with zero mean, then this estimation is precisely equivalent to the STA method. However, for non-Gaussian noise or spatial-temporally correlated input, STA provides biased and inaccurate estimates of RF structure. Here, following [17], we estimate RFs by minimising the quadratic approximation to the mutual information (QMI), without the Gaussian assumption, to provide a more accurate estimation of the mean of the receptive field parameters.

To the resulting RF estimates we further fitted a spatial-temporal Difference-of-Gaussian (ST-DoG) model to the receptive fields estimated from QMI method. This process reduced the dimension of the receptive field parameters down to several spatial-temporal Gaussian components, for further analysis. Observing a bimodal distribution of an ON-OFF index parameter, we classified neurons into two ON and OFF classes, based on this property. This procedure revealed an asymmetries in several properties between the ON and OFF visual pathways.

## II. MATERIALS AND METHODS

### A. Experimental Methods

Experimental methods are described in detail in [18]. Briefly, mice (2-4 month old C57BL/6) were sedated with 0.5 mg/kg chlorprothixene and anaesthetised using 1-1.5% isoflurane in O_2_. Multi-electrode arrays (MEAs, Neuronexus A1×16-Poly2-5mm-50s-177-A1) were vertically inserted into the brain until robust visual responses to gratings or white/black squares at a depth of 2500-3200 *µ*m below the pial surface indicated that the dLGN had been found. Electrodes were coated in DiI (20 mg in 300 *µ*l DMSO) and recording locations verified with post-hoc histology. Visual stimuli were presented using custom-modified MATLAB scripts based on Psychtoolbox [19] for MATLAB (Math-Works). Several types of visual stimuli were presented, as described in [18]: contrast-modulated noise movies for RF mapping [20], as well as drifting grating stimuli of varying spatial frequency, temporal frequency, contrast and direction. Spike sorting was performed; basic analyses of this dataset are described in [18].

### B. Quadratic Mutual Information

The idea of determining the receptive field by maximising the mutual information between stimulus and response (spike train) was popularised by Sharpee [21], as this allows the receptive field to be estimated using a spatially and/or temporally correlated stimulus. Quadratic Mutual Information (QMI) was proposed by Kapur as an alternative to the full mutual information measure [22]. The use of QMI for dimensionality reduction was proposed by Torkkola et al. [23], as this is computationally cheaper to use as the gain (or inverse cost) function. These two ideas were recently combined for receptive field extraction [17]. The mutual information between two continuous random variables *S* and *R* can be written as

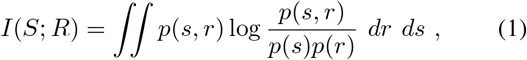

where, for instance, *S* might describe the visual stimulus, and *R* the neural response. In practice, as the stimulus space is extremely large, we might replace *S* by some transformed variable *Y*. The mutual information can be approximated by the quadratic form of mutual information [22] as

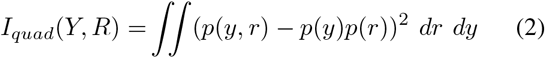

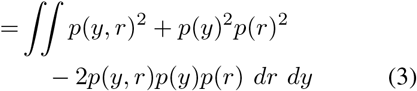

This latter quantity is much easier for calculation when the probability distributions are constructed from the data using kernel density estimation and the Gaussian kernel, because the integrals can be calculated in analytical form. The optimisation process:

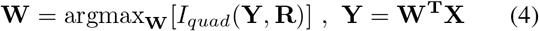

is unaffected, since the objective is to find the transformation **W** such that the mutual information between transformed inputs **Y** and weighted class labels **R** is maximised. For example the video input (S) is transformed into abstract lower dimensional feature (Y) and the quadratic mutual information between feature (Y) and cell spiking (R) is maximised.

Spike trains were binned by stimulus frames, with a duration of 50 ms (effective frame rate 20 Hz, although the monitor refresh rate was 120 Hz). The number of spikes within each frame bin is denoted by {*r*_*t*_}_*t*=1:*T*_. We extracted the aligned stimulus window *s*_(*t*−*τ*:*t*)_ for each bin, up to a total window length of *τ* frames. The filtered signal {*y*_*t*_ := ⟨*W, s*_(*t*−*τ*:*t*)_ ⟩} _*t*=1:*T*_ is used to calculate the QMI with{*r*_*t*_} _*t*=1:*T*_, where *W* is the linear receptive field vector of the cell, and ⟨.,. ⟩ is the inner product.

The original quadratic mutual information method was implemented by Katz et al. [17] for CPU in Matlab, but takes a long time to run (30-45 minutes per cell on our computer). We thus modified the method to incorporate GPU acceleration (with CUDA support), significantly reducing the computation time per cell (to 30-45 seconds per cell). Because the QMI method involves the manual setting of a bandwidth parameter *b*, Katz et al iteratively searched by first increasing then decreasing the bandwidth to find the suitable bandwidth value. In addition, they used the Matlab function *fminsearch* to find the optimal step-wise learning rate *γ* and then used this learning rate to update the receptive field vector *W* for each bandwidth chosen. The pseudo-code for the algorithm is shown below. We instead used *fminucn* with default *quasi-newton* optimization, finding a slight speed improvement. As bandwidth selection for kernel density estimation is an extensive research topic on its own, we do not go into detail here. Pseudocode for the algorithm is shown in Algorithm 1.

#### Algorithm 1: Pseudocode for the quadratic mutual information algorithm

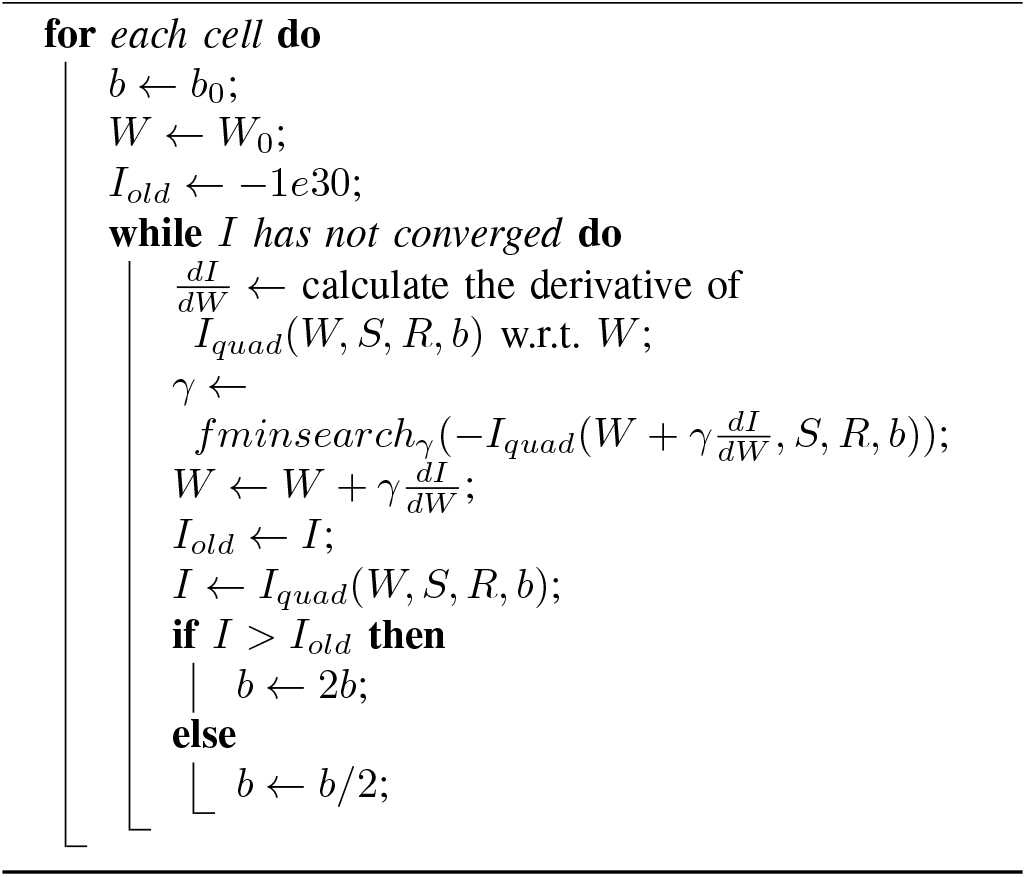

### C. RF analysis with the Spatial-temporal Difference of Gaussians model

We fit a spatial 2-dimensional and temporal 1-dimensional difference of Gaussians model to all receptive field vectors *W* to reduce the number of parameters and make easier further clustering and analysis.

The original Difference-of-Gaussian (DoG) model consists of two spherical spatial Gaussian components with identical center position, whereas the amplitude of the two spatial Gaussian components follows reversal exponential. However, we found that the standard model cannot capture all spatiotemporal properties of the receptive field vectors we obtained from recordings from the mouse dLGN. In particular, many receptive fields estimated have a non-spherical shape, i.e. a non-diagonal sigma for spatial Gaussian components. In addition, two spatial Gaussian components often do not have the same peak center in our dataset. Finally, the model needed to be modified to allow for a larger magnitude (up to complete) temporal reversal in amplitude.

We have thus loosened the assumption and allowed the center and surround Gaussian component to have different peak position. The Gaussian component has also been modified to allow an elliptic spatial shape. In addition, instead of a weak reversal, we assumed a difference of two Gaussians for each amplitude in the temporal domain. The new model with 16 parameters, which we term the Spatial-Temporal Difference of Gaussians model (ST-DoG), is defined in full in the Appendix, together with model comparison showing that it provides a better fit to RF parameters than the traditional DoG model.

### D. ON-OFF Classification

To classify cells into ON and OFF classes, we used the amplitudes of the ON and OFF components from the ST-DoG model fit, we define the ON-OFF index as:

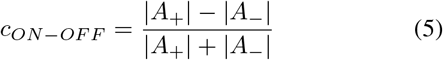

where *A*_+_ is the ON and *A*_−_ the OFF component amplitude parameter respectively. A value of *c*_*ON*−*OFF*_ = 1 of this index corresponds to an ON-only cell, and *c*_*ON*−*OFF*_ = − 1 corresponds to an OFF-only cell; cells with values lying in between these values possess both ON and OFF components. We use the value 0 as the decision boundary, i.e. if the ON-OFF index is less than 0, the cell is classified as an OFF cell. For example, in Fig 1 the *A*_+_ is the amplitude/peak of the yellow ST-Gaussian component, and similarly for *A*_−_ with blue ST-Gaussian component (for details of these parameters, refer to Appendix A).

**Fig 1.**
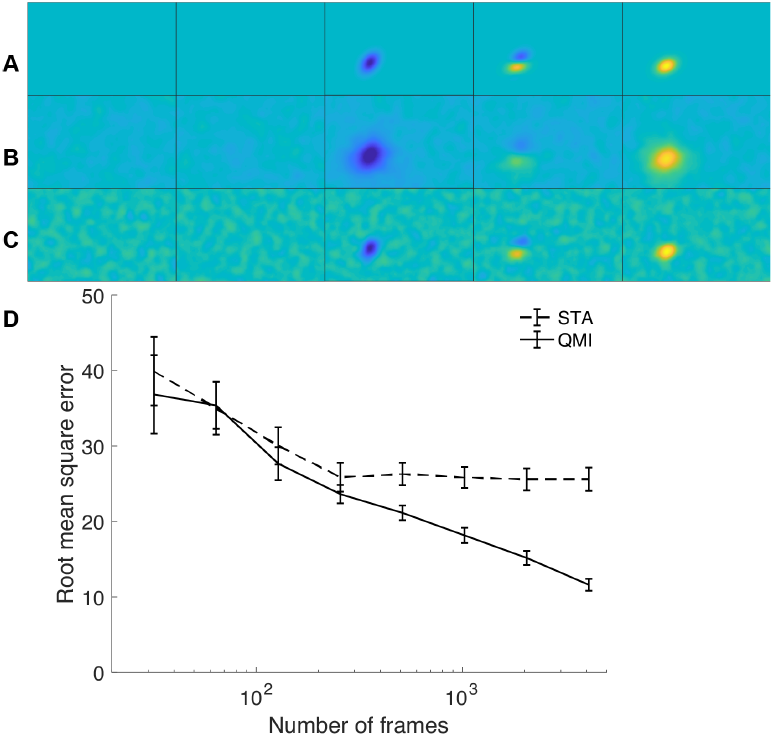
Comparison of STA and QMI reconstructions for a simulated receptive field (RF). (a) “Ground truth” receptive field (constructed by us, but based on one of the cells in the dataset) used to simulate spike trains using a linear-nonlinear-Poisson neuron model. 50 ms frames shown across horizontally, of size 105 × 65 degrees. (b) Spike-triggered average (STA) reconstruction of the RF for 6000 frames. (c) QMI reconstruction of the RF for 6000 frames. (d) RMS error for increasing number of spikes used in the estimation (averaged over 10 different receptive fields) for STA and QMI RF estimates, 12 by 12 visual degree around the center of the RF was been used for calculation.

## III. RESULTS

We first analysed the performance of the QMI algorithm for mapping synthetic receptive fields generated by simulating spike trains using a linear-nonlinear Poisson model, with linear filters derived from actual RFs observed in the dataset. An example is shown in Fig. 1a, depicting a “ground truth” RF over five 50 ms frames prior to spiking (spike occurrence in time at right of figure). In this example STA (Fig. 1b) over-estimates the size of the RF, which we assume is due to the biasing effect of spatial correlations in the contrast-modulated noise movie stimulus. In contrast, QMI more accurately estimates the receptive field (Fig. 1c), although apparently with the penalty of slightly more ripple in the baseline. This is an insubstantial issue, however, as the noise floor below a threshold can be trivially set to zero. QMI performance exceeds that of STA for datasets comprising 300 or more stimulus frames (Fig. 1d), or 15 seconds of data. For comparison, many minutes of data are normally collected in order to map RFs.

In the original analysis of [18], 189 cells were assessed as both having good quality spike sorting and an adequate receptive field. Using QMI RF mapping we were able to increase this to 285. We believe that this is primarily due to the use of STA for RF mapping in [18], with the consequence of sampling limitations being the need average the RFs across time, i.e. no attempt was made to extract the temporal component of the RF. While this produces an adequate spatial RF for cells strongly dominated by ON or OFF components, or with ON and OFF components largely spatially non-overlapping, it appears that there are a significant number of cells in the mouse dLGN with spatially overlapping ON and OFF components that come close to cancelling out if integrated over time to produce a single spatial RF. Several examples can be seen in Fig. 2.

**Fig 2.**
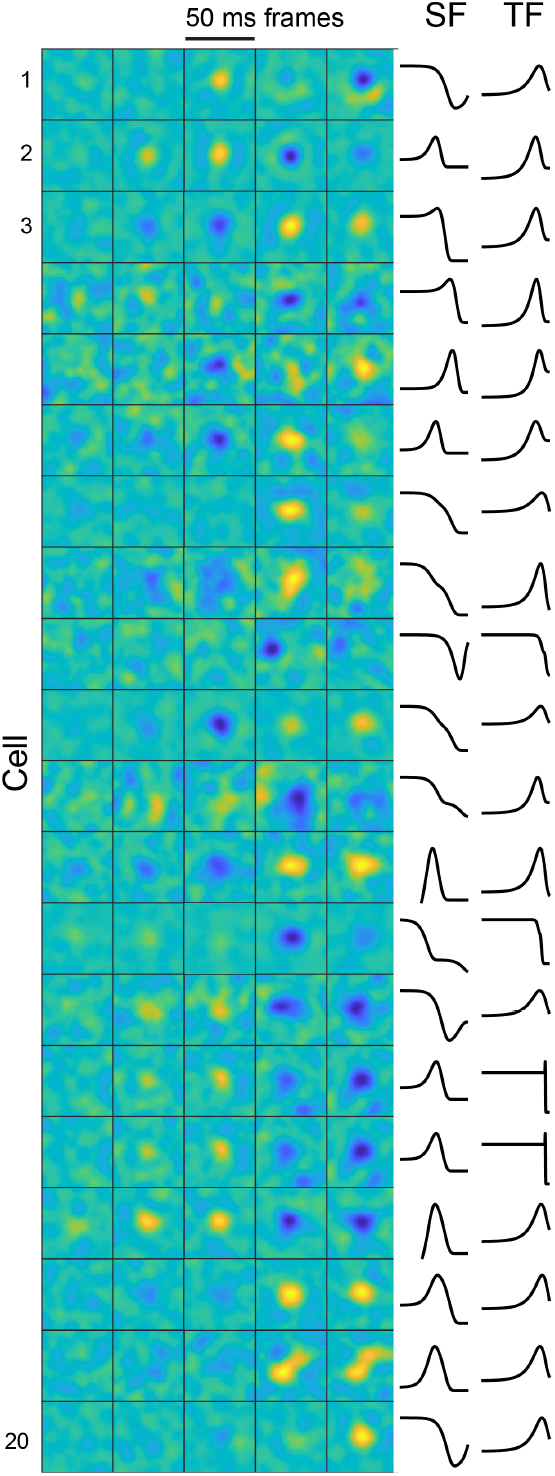
Example RFs obtained by QMI optimization. Each square shows a 20 × 20 degree area centred around the RF; 5 temporal frames of the RF are shown, with the far right being the first prior to spike onset. At right are shown the spatial (SF) and temporal frequency (TF) tuning curves (fit as in [18]) for each of the 20 cells. SFs range logarithmically from 0.01 to 0.96 cyc/deg, and TFs from 0.1 to 9.6 cyc/sec.

Of these cells, 133 were classified as ON and 152 as OFF, i.e. 46.6% ON to 53.4% OFF. Principal Component Analysis (Fig. 3) supported our conclusion that there were two broad categories of cells with differing properties, with nevertheless a region of parameter space where the two categories meet and are not (apart from the sign of the ON-OFF index) distinguishable. We observed a bi-modal distribution of ON-OFF indices (Fig. 4a), despite the continuous distribution ranging between −1 and 1.

**Fig 3.**
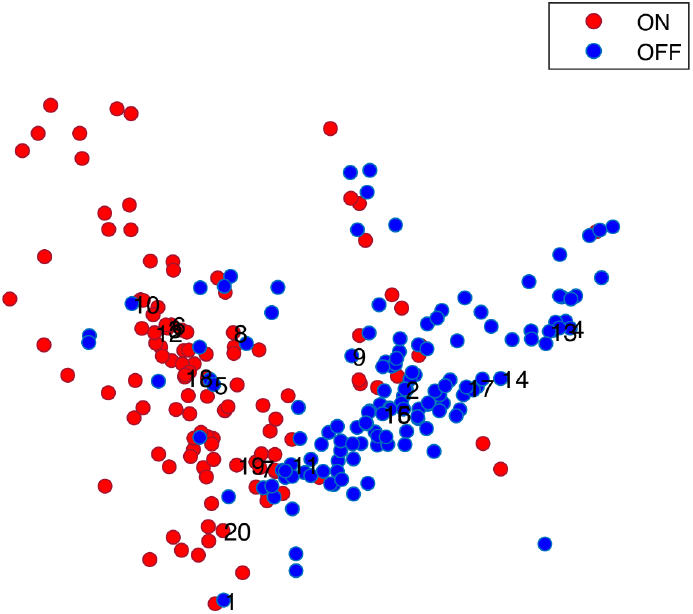
Principal Component Analysis (PCA) of receptive field structure. PCA was performed to reduce the dimensionality of the description of the RF structure extracted from the ST-DoG model from 16 to 2 parameters for each cell. Numbers reflect cells shown in Fig. 2.

**Fig 4.**
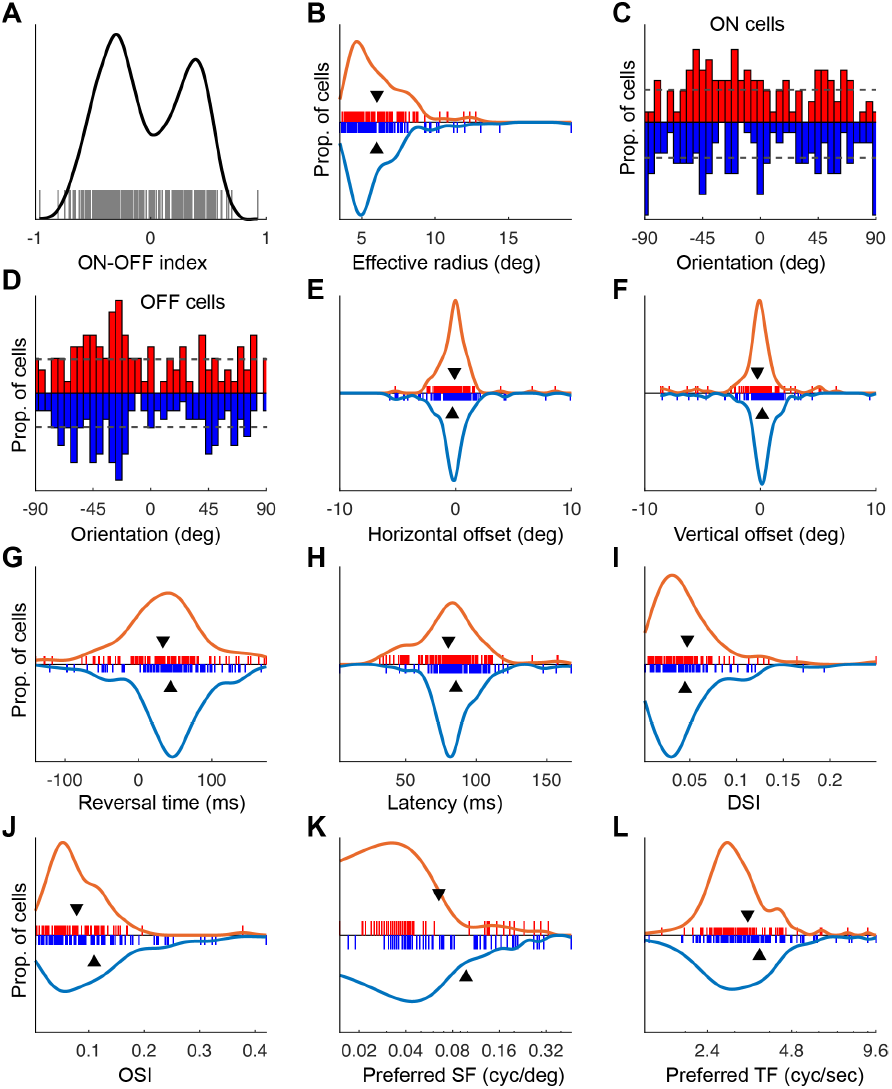
Analysis of the functional properties of dLGN ON and OFF receptive fields. (a) The distribution of ON-OFF indices for all 285 cells. (b) Effective radius for ON (red) and OFF (blue) cells. (c) Orientation (from − *π/*2 to *π/*2) of the ON component (red) and OFF component (blue) of the RF, for ON cells only. (d) As in (c), but for OFF cells. (e,f) Horizontal and vertical offset of ON and OFF RF components respectively, for ON cells (red) and OFF cells (blue). (g, h) Reversal time and latency for ON (red) and OFF cells (blue). (i) Direction selectivity Index (DSI). (j) Orientation Selectivity Index (OSI). (k) Preferred spatial frequency. (l) Preferred temporal frequency.

We extracted structural and functional properties of the RFs from the ST-DoG model. The effective RF radius, defined as the square root of the sum of the two “variances” of the Gaussian components 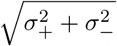 (see Appendix) did not differ substantially between ON and OFF cells (Fig. 4b). The ST-DoG model yields additional structural parameters, including the orientation of ON and OFF *components* of the receptive fields, shown for ON cells in Fig. 4c and OFF cells in Fig. 4d. There was no obvious asymmetry in these orientation parameters. Similarly, no difference between ON and OFF cells was apparent in the horizontal (*h*_+_ − *h*_−_) or vertical (*v*_+_ − *v*_−_) offsets between ON and OFF components (Fig. 4e and f). An additional temporal structural parameter is the reversal time, or the time between the two temporal Gaussian peaks, *τ*_+_ − *τ*_−_ for ON cells and *τ*_−_ − *τ*_+_ for OFF cells(mean 43.768 *±* 4.662 for OFF cells versus 33.415 *±* 5.727 for ON cells; p=0.159, two-sided t-test, n=133 ON and 152 OFF cells). ON cells showed on average a slightly (but not significantly) shorter latency, defined as *min* {*τ*_+_, *τ*_−_ }, the time to reach the first peak of either component, than OFF cells (mean 0.169 ± 0.215 (s.e.m.), p=0.182, two-sided t-test, n=133 ON and 152 OFF cells).

We were able to relate the ST-DoG parameters to functional measurements obtained from tuning curve mapping experiments using drifting gratings. The Direction Selectivity Index (see [18], measured by ∑ *F* (*θ*)*e*^2*iθ*^*/* ∑ *F* (*θ*) where *F* (*θ*) is the direction tuning curve) was similar for both ON and OFF cells (p=0.681, two-sided t-test, n=95 and 98 respectively), Fig. 4i. This was unsurprising given the relatively low number of direction selective cells within this dataset. OFF cells, however, were more orientation sensitive (Orientation Selectivity Index given by ∑ *F* (*θ*)*e*^*iθ*^*/* ∑ *F* (*θ*); mean 0.1099 ± 0.00882 for OFF cells versus 0.0783 ± 0.00583 for ON cells; p=0.0036, two-sided t-test, n=95 and 98). OFF cells were similarly on average selective for higher spatial frequencies (mean 0.0978 ± 0.0107 for OFF cells versus 0.0653 ± 0.0094 for ON cells; p=0.0256, two-sided t-test, n=142 OFF cells and 130 ON cells). OFF cells were sensitive to slightly higher temporal frequencies (mean 3.683 ± 0.1389 for OFF cells versus 3.342 ± 0.110 for ON cells; p=0.0644, two-sided t-test, n=140 OFF cells and 124 ON cells). Sample sizes differ in the above analyses because of the presence of low-pass tuned cells for whom a preferred spatial or temporal frequency was not defined.

## IV. DISCUSSION

We used quadratic mutual information to map RFs in the mouse dLGN. This yielded accurate, minimally biased estimates with substantially fewer samples than required by the method in most common use, the spike-triggered average. While it is well appreciated that the use of natural movie stimuli for receptive field mapping using spike-triggered approaches will result in biased RF estimates without the application of a correction procedure [24], such corrections can amplify noise and thus have adverse effects [25]. Information maximisation methods [21], [26] provide a much better approach for real-world stimuli, however the Kullback-Leibler mutual information is difficult to calculate and optimise, particularly for limited samples, meaning that approximations must usually be made. One such approximation, used in [21], is the use of the first order approximation to the information per spike [27]; this assumes independence of spikes, which is valid for e.g. the Poisson simulation we have used for validation, but does not hold for real world spike trains. An alternative approach, which we use here, is to use the Quadratic Mutual Information approximation [22]. As well as being easy to compute, this is differentiable, making it amenable to gradient-based optimization. In addition to advantages with respect to bias, QMI mapping of RFs is substantially more statistically efficient than STA mapping, requiring less recording duration to obtain RF structure.

QMI RF mapping yielded substantially more information about RF structure on an existing dataset [18], allowing the RFs of more cells to be obtained (285 as opposed to 189), and providing space-time RFs rather than purely spatial RFs that averaged over time. This allowed additional insight to be gained, and the enhanced dataset, while essentially yielding a picture of structural symmetry between ON and OFF receptive fields, did reveal asymmetries in a number of functional properties between the ON and OFF pathways in the mouse dLGN, including the orientation selectivity index and preferred spatial frequency (with a weaker effect apparent for the preferred temporal frequency). This implies some degree of functional specialisation of the ON and OFF pathways. While the role of this pathway splitting remains speculative (see e.g. [28]): one reason may relate to differences in uncertainty in dark versus well-illuminated areas of a visual scene.

One limitation of the dataset analysed was the use of a 20 Hz refresh rate for frame updates. This limited the temporal fidelity with which temporal aspects of RF structure can be extracted. Given that the QMI method enables statistically efficient extraction of spatiotemporal RF structure with high temporal fidelity, it will be of interest to collect additional experimental data with higher stimulus refresh rates. Analysis of finer temporal structure of the spatiotemporal RFs of dLGN neurons may reveal additional diversity in information processing functionality [29]. In addition, further analysis can be done by comparing this with other methods such as variants of spike-triggered covariance for highly correlated stimuli. However, the use of high dimensional visual stimuli can make this computationally expensive.

We finally remark that, although here, the QMI receptive field mapping method has been applied to an electrophysiological dataset, it is likely to have wider applications. In particular, we expect it to find use in mapping sensory receptive fields through technology such as multiphoton calcium [30], [31] or voltage [32] imaging, for studying other mammals with higher visual acuity, and in additional domains such as mapping selectivity functions in spatial memory [33].

## APPENDIX

In this appendix we describe the ST-DoG model, and compare its ability to account for the diversity of RF properties with the standard DoG model. The new model has the following parameters: *h*_+_, horizontal position of the positive Gaussian components; *v*_+_, vertical position of the positive Gaussian components; *h*_−_, *v*_−_, similar for the negative Gaussian components; *σ*_*h*+_,*σ*_*v*+_,*θ*_+_, eclipse radius and angle of the positive spatial Gaussian component; *σ*_*h*−_,*σ*_*v*−_,*θ*_−_, eclipse radius and angle of the negative spatial Gaussian component; *A*_+_, *τ*_+_, *σ*_+_, the temporal positive Gaussian component; *A*_−_, *τ*_−_, *σ*_−_, the temporal negative Gaussian component. We first define:

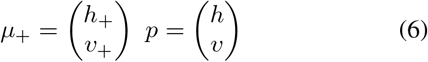

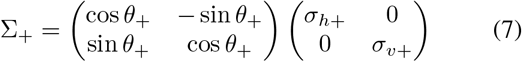

And similarly for *μ*_−_ and ∑_−_. The receptive field is then defined with those parameters by:

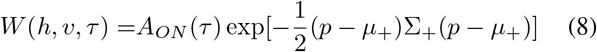

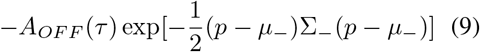

Amplitude for ON components (similar for OFF):

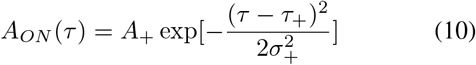

We restricted *A*_+_ and *A*_−_ to be positive and used a non-linear optimisation algorithm to find the optimal parameters (fmincon in Matlab).

Figure 5 indicates that the ST-DoG model (16 parameters) provides a better account than the DoG model with 10 parameters. Note that the error is capped due to use of constrained non-linear optimization, and that a small group of neurons were fitted better by the DoG model. Considering model selection (Akaike Information Criterion, AIC), the number of data points for the 10×10 degree window around the RF centre for curve fitting is much larger than the difference in parameter count between the models, thus model error is the dominant AIC term, and the ST-DoG model provides a better account of RF properties.

**Fig 5.**
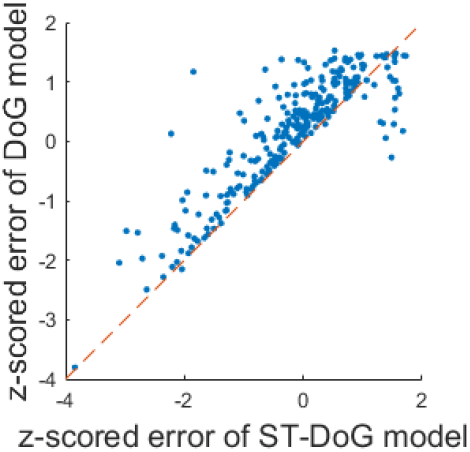
Comparison of the ST-DoG and DoG models.

## References

[1] H. von Helmholtz, “Handbuch der physiologischen optik / Handbook of Physiological Optics (J. P. C. SouthallTrans. 3rd ed.),” Allgemeine Encyklopadie der Physik Vol 2, ed Karsten G(Voss, Leipzig, Germany)/Dover, pp. 186–193, 1867/(1910, 1962).

[2] G. Galilei, “Dialogue concerning the two chief world systems,” Battista Landini, Florence, Italy, 1632.

[3] P. H. Schiller, “Central connections of the retinal on and off pathways,” Nature, vol. 297, pp. 580–583, 1982.

[4] P. H. Schiller, J. H. Sandell, and J. H. R. Maunsell, “Functions of the on and off channels of the visual system,” Nature, vol. 322, pp. 824–825, 1986.

[5] F. M. de Monasterio, “Asymmetry of on- and off-pathways of blue-sensitive cones of the retina of macaques,” Brain Research, vol. 166, no. 1, pp. 39–48, 1979.

[6] J. Haag, A. Mishra, and A. Borst, “A common directional tuning mechanism of Drosophila motion-sensing neurons in the on and in the off pathway,” eLife, p. e29044, 2017.

[7] K. A. Zaghloul, K. Boahen, and J. B. Demb, “Different circuits for on and off retinal ganglion cells cause different contrast sensitivities,” Journal of Neuroscience, vol. 23, no. 7, pp. 2645–2654, 2003.

[8] J. Veit, A. Bhattacharyya, R. Kretz, and G. Rainer, “On the relation between receptive field structure and stimulus selectivity in the tree shrew primary visual cortex,” Cereb. Cortex, vol. 24, no. 10, pp. 2761– 2771, 2014.

[9] D. I Vaney, B. Sivyer, and W. Taylor, “Direction selectivity in the retina: Symmetry and asymmetry in structure and function,” Nature Reviews Neuroscience, vol. 13, pp. 194–208, 02 2012.

[10] J. Z. Jin, C. Weng, C.-I. Yeh, J. A. Gordon, E. S. Ruthazer, M. P. Stryker, H. A. Swadlow, and J.-M. Alonso, “On and off domains of geniculate afferents in cat primary visual cortex,” Nat. Neurosci., vol. 11, no. 1, p. 88, Dec 2007.

[11] D. Xing, C.-I. Yeh, J. Gordon, and R. M. Shapley, “Cortical brightness adaptation when darkness and brightness produce different dynamical states in the visual cortex,” Proc. Natl. Acad. Sci. U.S.A., vol. 111, no. 3, pp. 1210–1215, Jan 2014.

[12] D. A. Clark, J. E. Fitzgerald, J. M. Ales, D. M. Gohl, M. A. Silies, A. M. Norcia, and T. R. Clandinin, “Flies and humans share a motion estimation strategy that exploits natural scene statistics,” Nature Neuroscience, vol. 17, no. 2, pp. 296–303, 2014.

[13] K. M. Ahmad, K. Klug, S. Herr, P. Sterling, and S. Schein, “Cell density ratios in a foveal patch in macaque retina,” Visual Neuroscience, vol. 20, no. 2, p. 189, 2003.

[14] S. Ravi, D. Ahn, M. Greschner, E. Chichilnisky, and G. D. Field, “Pathway-specific asymmetries between on and off visual signals,” Journal of Neuroscience, vol. 38, no. 45, pp. 9728–9740, 2018.

[15] V. Zemon, J. Gordon, and J. Welch, “Asymmetries in on and off visual pathways of humans revealed using contrast-evoked cortical potentials,” Visual Neuroscience, vol. 1, no. 1, p. 145–150, 1988.

[16] A. Leonhardt, G. Ammer, M. Meier, E. Serbe, A. Bahl, and A. Borst, “Asymmetry of drosophila on and off motion detectors enhances real-world velocity estimation,” Nature Neuroscience, vol. 19, 02 2016.

[17] M. L. Katz, T. J. Viney, and K. Nikolic, “Receptive field vectors of genetically-identified retinal ganglion cells reveal cell-type-dependent visual functions,” PLOS ONE, vol. 11, no. 2, pp. 1–29, 2016.

[18] J. Tang, S. C. Ardila Jimenez, S. Chakraborty, and S. R. Schultz, “Visual receptive field properties of neurons in the mouse lateral geniculate nucleus,” PLOS ONE, vol. 11, no. 1, pp. 1–34, 01 2016.

[19] M. Kleiner, D. Brainard, and D. Pelli, “What’s new in Psychtoolbox-3?” Perception, vol. 36 EVCP Abstract Supplement, 2007.

[20] C. M. Niell and M. P. Stryker, “Highly selective receptive fields in mouse visual cortex,” Journal of Neuroscience, vol. 28, no. 30, pp. 7520–7536, 2008.

[21] T. Sharpee, “Neural Responses To Natural Signals: Maximally Informative Dimensions,” Neural Computation, vol. 16, pp. 223–250, 2004.

[22] J. Kapur, Measures of Information and Their Applications. Wiley, 1994.

[23] K. Torkkola, “Feature extraction by non-parametric mutual information maximization,” Journal of Machine Learning Research, vol. 3, pp. 1415–1438, 2003.

[24] J. Touryan, G. Felsen, and Y. Dan, “Spatial structure of complex cell receptive fields measured with natural images,” Neuron, vol. 45, no. 5, pp. 781–791, 2005.

[25] F. E. Theunissen, K. Sen, and A. J. Doupe, “Spectral-temporal receptive fields of nonlinear auditory neurons obtained using natural sounds,” Journal of Neuroscience, vol. 20, no. 6, pp. 2315–2331, 2000.

[26] T. O. Sharpee, “Comparison of information and variance maximization strategies for characterizing neural feature selectivity,” Statistics in medicine, vol. 26, no. 21, pp. 4009–4031, 2007.

[27] W. Skaggs, B. Mcnaughton, and K. Gothard, “An information-theoretic approach to deciphering the hippocampal code,” Advances in neural information processing systems, vol. 5, pp. 1030–1037, 1992.

[28] J. Gjorgjieva, H. Sompolinsky, and M. Meister, “Benefits of pathway splitting in sensory coding,” Journal of Neuroscience, vol. 34, no. 36, pp. 12 127–12 144, 2014.

[29] D. M. Piscopo, R. N. El-Danaf, A. D. Huberman, and C. M. Niell, “Diverse Visual Features Encoded in Mouse Lateral Geniculate Nucleus,” Journal of Neuroscience, vol. 33, no. 11, pp. 4642–4656, 2013.

[30] S. L. Smith andM. Häusser, “Parallel processing of visual space by neighboring neurons in mouse visual cortex,” Nature neuroscience, vol. 13, no. 9, p. 1144, 2010.

[31] S. R. Schultz, C. S. Copeland, A. J. Foust, P. Quicke, and R. Schuck, “Advances in two-photon scanning and scanless microscopy technologies for functional neural circuit imaging,” Proceedings of the IEEE, vol. 105, no. 1, pp. 139–157, 2016.

[32] P. Quicke, C. L. Howe, P. Song, H. V. Jadan, C. Song, T. Knöpfel, M. Neil, P. L. Dragotti, S. R. Schultz, and A. J. Foust, “Subcellular resolution three-dimensional light-field imaging with genetically encoded voltage indicators,” Neurophotonics, vol. 7, no. 3, p. 035006, 2020.

[33] M. A. Go, J. Rogers, G. P. Gava, C. Davey, S. Prado, Y. Liu, and S. R. Schultz, “Place cells in head-fixed mice navigating a floating real-world environment,” Frontiers in Cellular Neuroscience, p. 10.3389/fncel.2021.618658, 2021.

